# Dynamical Interrogation of Serpentine Medulla Circuits in the *Drosophila* Optic Lobe

**DOI:** 10.64898/2026.01.08.698345

**Authors:** Nalin Dhiman, Siddharth Panwar

## Abstract

Dense connectomes invite a tempting leap: if we can “wire up” a circuit, perhaps its dynamics (and even its function) will emerge. In practice, connectomic constraints leave large biophysical degrees of freedom (synapse signs/strengths, neuromodulation, gap junctions), so qualitative dynamical claims can be fragile. We present a simple *forensic* workflow for testing connectome-to-dynamics hypotheses that is deliberately hostile to wishful thinking: each stage is gated by sanity checks, run across multiple random seeds, and reported with effect sizes and negative controls. As a case study, we instantiate the Serpentine Medulla (Sm) interneuron family from FlyWire as a current-based adaptive exponential integrate-and-fire (AdEx) network and test a sharp narrative: whether strong recurrence supports a working-memory-style bistable attractor. Under strict attractor criteria (noise removed after the kick), we find no robust evidence for bistability (paired Wilcoxon *p* = 0.099, within-subject *d*_*z*_ = − 0.25). For oscillations, a corrected reproduction across 20 random seeds shows that recurrent coupling strongly *amplifies* integrated 30–80 Hz power when comparing the fully coupled condition (Full *W*) to an uncoupled control (*W* = 0) (paired Wilcoxon *p* = 1.9 × 10^−6^; *d*_*z*_ = 5.96). We also observe a fast negative-coupling signature in input decomposition (mean peak correlation *r* ≈ − 0.40 at ∼ 4 ms lag), while stressing that these signatures are conditional on the chosen model class and operating point. Overall, the main contribution is methodological: an auditable template for *falsifying* attractive connectome-based stories rather than polishing them.

## 1 Introduction

### From wiring diagrams to dynamics: a deliberately conservative stance

Cellular-resolution connectomes are arriving faster than our ability to interpret them. FlyWire provides a brain-wide synapse-level map of adult *Drosophila* circuitry with community-driven proofreading [1]. In parallel, optic-lobe annotation has matured into large-scale cell-type catalogs and wiring diagrams that organize early visual circuits into structured families [2, 3]. These datasets make it tempting to “simulate the connectome” and read off function.

But a connectome is not a dynamical model. Connectomics constrains *who* connects to *whom* (and approximately how strongly via synapse counts), but typically leaves unresolved: (1) the *sign* and effective strength of synapses (including state dependence); (2) intrinsic excitability distributions across cell types; and (3) non-synaptic interactions (neuromodulators, electrical coupling, glia). As a result, the mapping from graph to dynamics is underdetermined: many dynamical systems are consistent with the same adjacency matrix. Recent work argues that connectomes can still guide mechanistic modeling, especially when paired with perturbation experiments and fitted dynamics [4]. Our goal here is more conservative: before optimizing high-capacity models, we ask whether *qualitative* dynamical claims survive adversarial falsification within a fixed, transparent model class.

### Biological context: the optic lobe, the medulla, and the Serpentine Medulla family

The fly optic lobe is a laminated, retinotopically organized set of neuropils (lamina, medulla, lobula, lobula plate) that performs the early transformations of visual signals, routing features into parallel pathways [5]. The medulla, in particular, is a layered neuropil that receives direct photoreceptor input (R7/R8) and lamina outputs (R1–R6 via lamina neurons), and contains a diverse population of columnar and tangential interneurons that mediate feature extraction, pooling, and routing [5, 6]. Connectomic reconstructions show that medulla circuits are both stereotyped and variable in synapse counts across neighboring columns [6], highlighting a key point: anatomy supplies *constraints*, not a unique dynamical regime.

Within this medulla ecosystem, the Serpentine Medulla (Sm) family is described in recent optic-lobe cell-type catalogs as a large and diverse interneuron family concentrated near the medulla’s serpentine layer (around layer M7), with characteristic stratification and extensive recurrence [2]. In the wiring diagram, Sm types communicate with multiple transmedullary (Tm/Mt) and boundary neuron classes, and are discussed as part of the circuitry involved in chromatic processing within the optic lobe’s color subsystem [2]. The combination of (i) large population size, (ii) dense recurrence, and (iii) cross-layer coupling makes Sm a natural testbed for “connectome-to-dynamics” narratives.

### A sharp dynamical question: does Sm recurrence imply bistable memory?

Recurrent networks can support multiple dynamical motifs: gain control, oscillations, winner-take-all states, and (in some parameter regimes) multistable attractors often invoked as a substrate for working memory. It is therefore plausible—but not guaranteed—that a strongly recurrent connectome subgraph might realize bistability “by default” when instantiated as a spiking network.

We treat this as a falsifiable hypothesis. Specifically, we ask whether an Sm network, instantiated with adaptive exponential integrate-and-fire neurons and synapse-count-based coupling, supports a bistable high-activity state that persists after a strong transient input is removed. A critical methodological nuance is that persistent activity can be faked by slow relaxation under ongoing noise. To avoid that trap, our attractor assay explicitly turns *off* stochastic drive after the kick and demands state separation in deterministic dynamics.

### Contribution: a “forensic” pipeline and a case study

The main contribution is methodological. We formalize a pipeline with: (i) nontriviality gates (avoid silent or saturated regimes), (ii) repeated trials across random seeds, (iii) explicit effect sizes (not just *p*-values), (iv) negative controls (simple linear dynamical baselines), and (v) stress tests for uncertain assumptions (e.g., neurotransmitter-to-sign mapping). We then apply it end-to-end to the Sm family as a case study, and report both positive and negative findings without functional overreach.

## 2 Methods

### 2.1 Connectome extraction and representation

We analyze the Serpentine Medulla (Sm) family extracted from the FlyWire connectome [1] using optic-lobe type labels from recent cell-type catalogs [2, 3]. The analysis focuses on the Sm × Sm subgraph (internal recurrence) with *N* = 4463 neurons (nodes). Edges are weighted by synapse counts.

Let *W* ∈ ℝ^*N×N*^ denote the synapse-count adjacency matrix, where *W*_*ij*_ ≥ 0 is the number of chemical synapses from presynaptic neuron *j* to postsynaptic neuron *i*. We convert synapse counts to signed current increments by assigning each edge a sign *σ*_*ij*_ ∈ {−1, +1} and scaling by a global factor *α >* 0 (details below).

### 2.2 Spiking network model (AdEx + exponential synapses)

Each neuron *i* ∈ {1, …, *N*} is modeled as an adaptive exponential integrate-and-fire (AdEx) unit [7]. The membrane potential *v*_*i*_(*t*) and adaptation variable *w*_*i*_(*t*) evolve as

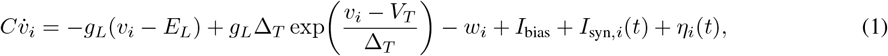

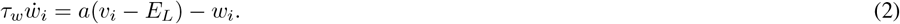

**Table 1:**
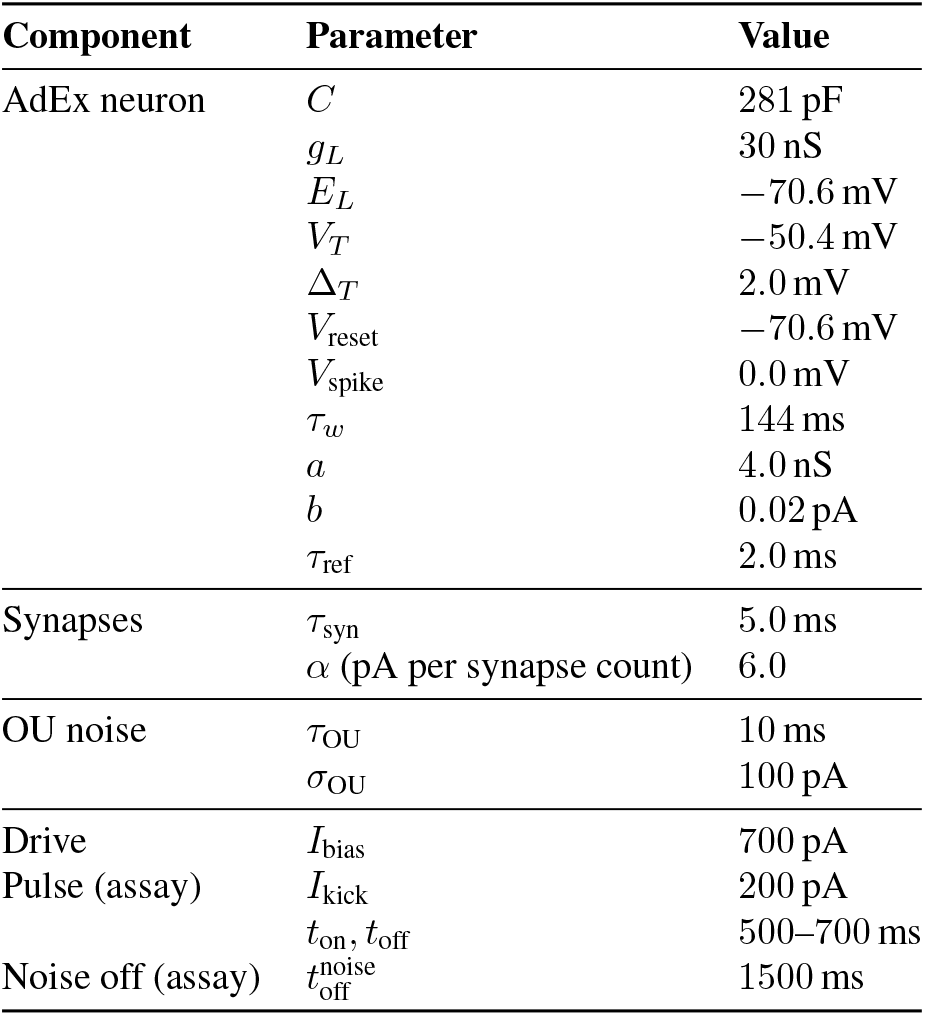
Default model parameters used in this work. Units are chosen so that 1 nS · 1 mV = 1 pA, consistent with Eqs. (1)–(3).

| Component | Parameter | Value |
| --- | --- | --- |
| AdEx neuron | $C$ | 281 pF |
| | $g_L$ | 30 nS |
| | $E_L$ | −70.6 mV |
| | $V_T$ | −50.4 mV |
| | $\Delta_T$ | 2.0 mV |
| | $V_{\text{reset}}$ | −70.6 mV |
| | $V_{\text{spike}}$ | 0.0 mV |
| | $\tau_w$ | 144 ms |
| | $a$ | 4.0 nS |
| | $b$ | 0.02 pA |
| | $\tau_{\text{ref}}$ | 2.0 ms |
| Synapses | $\tau_{\text{syn}}$ | 5.0 ms |
| | $\alpha$ (pA per synapse count) | 6.0 |
| OU noise | $\tau_{\text{OU}}$ | 10 ms |
| | $\sigma_{\text{OU}}$ | 100 pA |
| Drive | $I_{\text{bias}}$ | 700 pA |
| Pulse (assay) | $I_{\text{kick}}$ | 200 pA |
| | $t_{\text{on}}, t_{\text{off}}$ | 500–700 ms |
| Noise off (assay) | $t_{\text{off}}^{\text{noise}}$ | 1500 ms |

When *v*_*i*_ crosses a spike threshold *V*_spike_, a spike is emitted and the state is reset: *v*_*i*_ ← *V*_reset_ and *w*_*i*_ ← *w*_*i*_ + *b*, followed by an absolute refractory period *τ*_ref_.

#### Current-based synapses

Synaptic input is implemented as an exponential filter driven by presynaptic spikes. Let *s*_*j*_(*t*) ∈ {0, 1} indicate whether neuron *j* spiked in the previous time bin. In vector form, with *s*(*t*) ∈ {0, 1}^*N*^, we update the synaptic current as

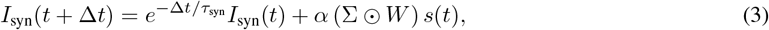

where *τ*_syn_ is a synaptic time constant and ∑ is the sign matrix with entries *σ*_*ij*_; ⊙ denotes elementwise multiplication.

#### Noise model (Ornstein–Uhlenbeck current)

To model fluctuating input and to avoid deterministic synchrony artifacts, each neuron receives independent Ornstein–Uhlenbeck (OU) current noise *η*_*i*_(*t*):

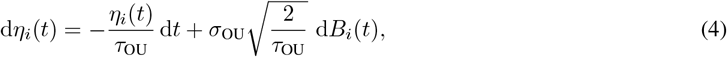

where *B*_*i*_(*t*) are independent standard Wiener processes. We implement the exact discrete-time update (Appendix C).

### 2.3 Parameterization and simulation protocol

Unless stated otherwise, simulations use the default parameters. These values are *not* claimed to be cell-type-specific fits; they define a transparent baseline operating regime.

#### Simulation timeline (used throughout)

Time is discretized with step Δ*t* = 0.1 ms for total duration *T* = 2000 ms. For the strict attractor assay, a uniform pulse current is applied from *t* = 500 to 700 ms and OU noise is set to zero after *t* = 1500 ms (deterministic tail).

### 2.4 Assumptions and calibration

#### Synapse sign policy and uncertainty

Synapse signs are assigned using FlyWire neurotransmitter predictions and a default mapping: acetylcholine → excitatory; GABA → inhibitory. Glutamate is treated as inhibitory by default, consistent with evidence for glutamatergic inhibition via glutamate-gated chloride channels in multiple *Drosophila* circuits [8, 9, 10]. Because neurotransmitter-based sign assignment is uncertain in general (and can depend on receptor expression), we treat sign policy as a target for stress testing. We perform robustness perturbations by randomly flipping a fraction *p* of nonzero edges (Section 2.5.1).

#### Global coupling scale calibration

Synapse counts provide relative coupling strengths but not absolute current amplitudes. We therefore introduce a global synapse-count-to-current scale *α* in Eq. (3). We sweep *α* and select a working point that produces: (i) bounded non-saturating firing rates; and (ii) nontrivial dynamics (not silent, not blown up). This is a pragmatic choice rather than a biological fit.

### 2.5 Assays and analysis metrics

#### Empirical transfer function and mean-field sanity checks

A useful “first check” is whether the chosen neuron model has a sensible input–output nonlinearity. We estimate an empirical transfer function *ϕ*(*I*) for the AdEx parameters by simulating a homogeneous population under constant input current *I* and measuring steady-state firing rate. We use *ϕ* for qualitative mean-field reasoning: a coarse mean-rate model for a recurrent population can be written as

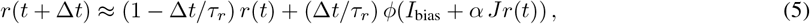

where *r*(*t*) is population rate and *J* is an effective coupling parameter (a strong simplification of the full heterogeneous network). In such models, bistability requires multiple stable fixed points, which in turn requires the gain (slope) of the right-hand side to exceed unity over some range. We use this only as a sanity lens, not a proof.

#### Bistability assay: strict “kick-and-noise-off” test

We test for bistability by comparing two conditions:

- **Control:** baseline simulation with OU noise.
- **Pulse:** an added transient “kick” current *I*_kick_ to all neurons for a window [*t*_0_, *t*_1_].

A key criterion is that after the pulse ends, we remove stochastic drive by setting *σ*_OU_ = 0 at time *t*_off_. If the system possesses two attractors, the kicked trajectory should remain separated from the control trajectory under deterministic dynamics. We quantify separation using the mean population firing rate over an end window [*T* − Δ, *T*] and compare control vs pulse across seeds using a paired Wilcoxon signed-rank test.

#### Population rate, spectral analysis, and gamma-band power

We define the instantaneous population spike count as *n*(*t*) = ∑_*i*_ *s*_*i*_(*t*) and the population rate as a smoothed version

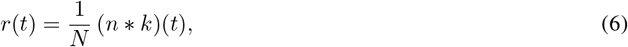

where *k* is a smoothing kernel (Gaussian or exponential; fixed across conditions). Power spectral densities (PSDs) are estimated with Welch’s method using a fixed windowing scheme across all comparisons. We define integrated gamma-band power as

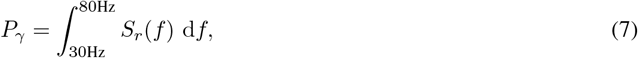

where *S*_*r*_(*f*) is the PSD of *r*(*t*). We compare *P*_*γ*_ across conditions using paired tests across seeds. Important caveat: a non-significant difference in *P*_*γ*_ should not be interpreted as evidence of equivalence; it may simply reflect limited sensitivity under the current sample size, noise level, and fixed preprocessing choices.

#### Input decomposition and external coupling signature

We decompose synaptic drive to a chosen subpopulation into recurrent and external contributions. Let 𝒮 denote the subpopulation (in our case, the Sm subcircuit under analysis), and let 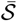denote the rest of the network in the embedded simulation. Then the synaptic current into 𝒮 can be written as

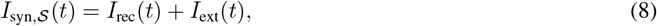

where *I*_rec_ arises from presynaptic spikes within 𝒮 and *I*_ext_ from spikes in 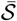. We compute the normalized cross-correlation between *r*_𝒮_ (*t*) and *I*_ext_(*t*) and report the most negative peak and its lag.

#### Linear dynamical baseline

To provide a negative control for “local predictive success”, we fit a one-dimensional linear dynamical system (LDS) to the population rate:

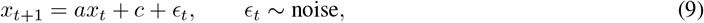

using ridge regression to stabilize the estimate. We evaluate: (i) *one-step prediction* under teacher forcing (using true *x*_*t*_), and (ii) *open-loop rollout* (feeding the model’s own predictions). A stable linear system generically collapses to a fixed point and cannot robustly represent a limit cycle unless tuned to the unit circle. This provides a caution: one-step prediction quality does not imply correct long-horizon dynamics.

#### 2.5.1 Stress test: random sign perturbations

We perturb the sign matrix Σ by flipping a random fraction *p* of nonzero edges. For each perturbation, we re-run the simulation and compute the mean absolute difference in population rate between baseline and perturbed conditions. This tests whether qualitative regime features depend on a knife-edge sign assignment.

### 2.6 Statistics and reproducibility

#### Statistics and reporting

Across-seed comparisons use paired Wilcoxon signed-rank tests. We report within-subject effect size *d*_*z*_:

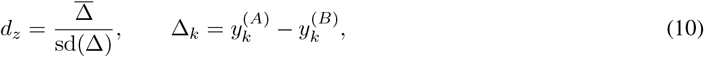

where *A* and *B* denote the two paired conditions under comparison (e.g., Pulse vs Control or Full *W* vs *W* = 0) and *k* indexes random seeds. Where relevant, we use bootstrap confidence intervals for descriptive quantities (Appendix C).

#### Correctness gates (“forensic checks”)

To prevent silent failure modes, scripts implement explicit checks including: rate bounds (avoid silent/saturated networks), monotonic transfer curves, fixed PSD parameters, transient removal for correlation estimates, and *p*-value range validation. Appendix A lists gates and their motivation.

## 3 Results

### 3.1 Calibration and baseline operating regime

#### Empirical AdEx transfer curve provides a calibrated nonlinearity

Figure 1 shows the empirically estimated transfer function *ϕ*(*I*) for the AdEx parameterization used in downstream simulations. The curve is monotone and saturating, consistent with the expected AdEx input–output behavior. We use the bias current *I*_bias_ (dashed line) to set a tonic operating regime with ongoing spiking.

**Figure 1:**
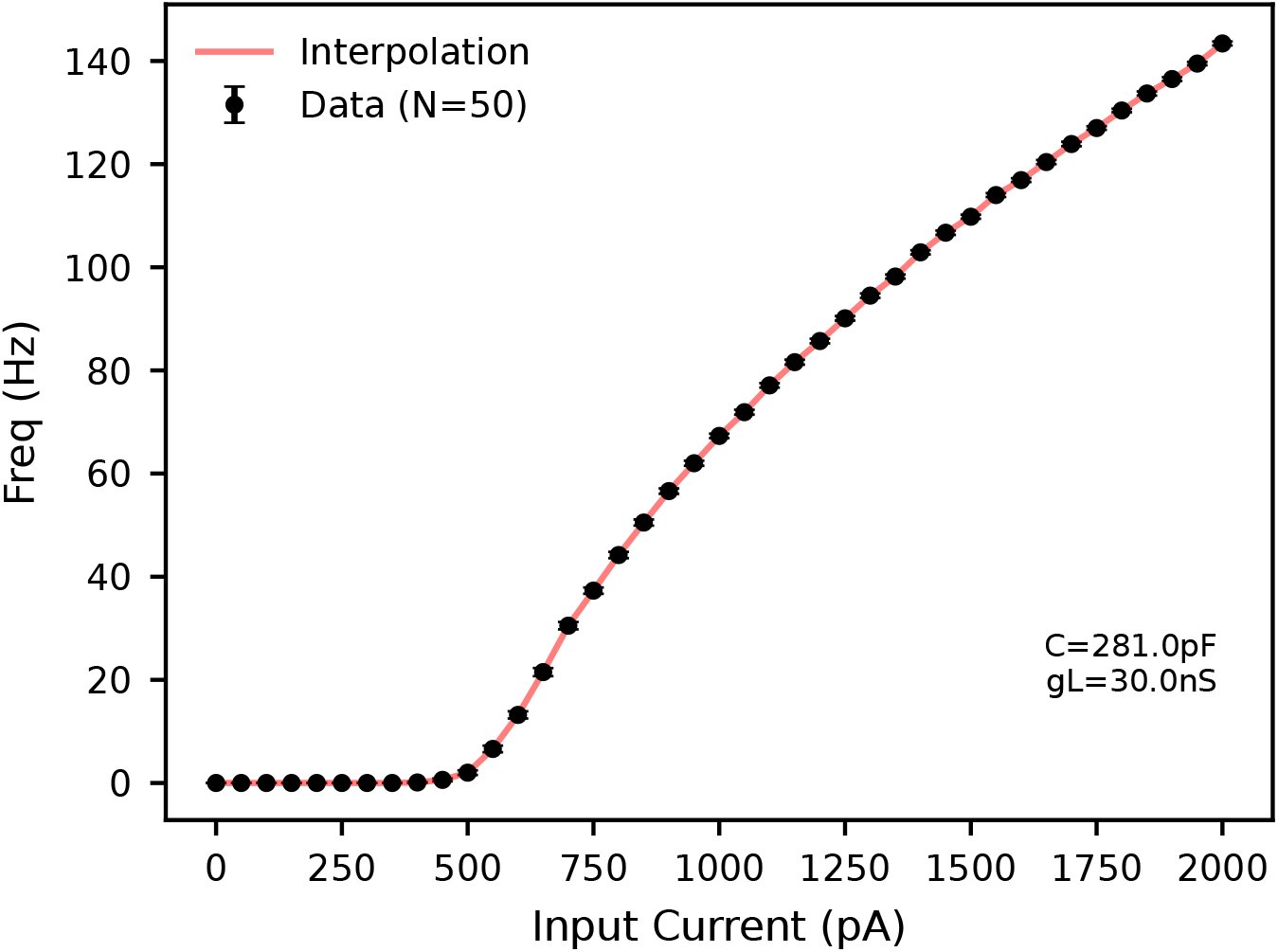
Empirical AdEx transfer function. Points show simulated steady-state firing rates for constant current drive; the curve is a smooth fit *ϕ*(*I*). The dashed line indicates the bias current used in the network simulations.

#### Sm connectivity exhibits structured sparsity and heavy-tailed degree

Figure 2A shows the sparsity pattern of the Sm *×* Sm adjacency matrix. Visual “bands” and heterogeneous density suggest non-random connectivity structure and subtype heterogeneity. Sorting by in-degree concentrates density into a core (Figure 2B), consistent with heavy-tailed degree. This structural heterogeneity matters dynamically: a homogeneous mean-field picture can miss hub-driven regimes.

**Figure 2:**
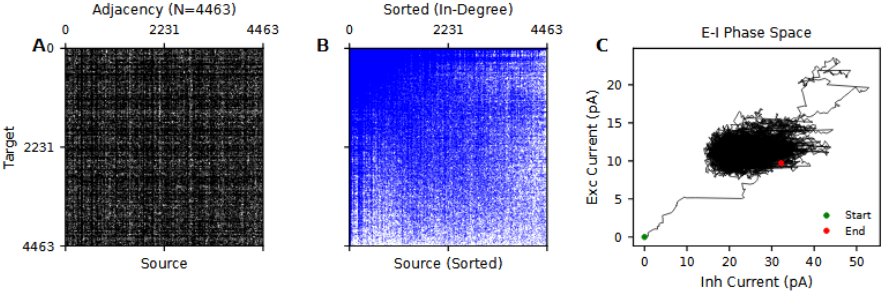
Sm connectivity and E–I operating regime. (A) Sparsity pattern of the Sm×Sm adjacency matrix (*N* = 4463). (B) Same matrix after sorting neurons by in-degree. (C) Mean excitatory synaptic current vs inhibitory current magnitude (seed 0), reconstructed from recorded spikes and signed weights using the same exponential synaptic kernel as the simulator.

We also reconstruct an excitatory–inhibitory (E–I) operating trajectory by splitting signed synaptic input into positive and negative components and applying the same synaptic filter used in the simulation. The trajectory rapidly leaves the quiescent origin and converges to a compact cloud in E–I space (Figure 2C), consistent with a stable ongoing regime.

#### Global coupling sweep selects a bounded, nontrivial regime

We sweep the global synaptic scaling *α* to select an operating point with nontrivial but bounded dynamics. Figure 3 shows mean population rate and a synchrony proxy (CV of smoothed population rate) versus *α*. We choose *α* = 6 as a working point balancing non-silence and non-runaway activity. This is a pragmatic calibration, not a fitted biophysical estimate.

**Figure 3:**
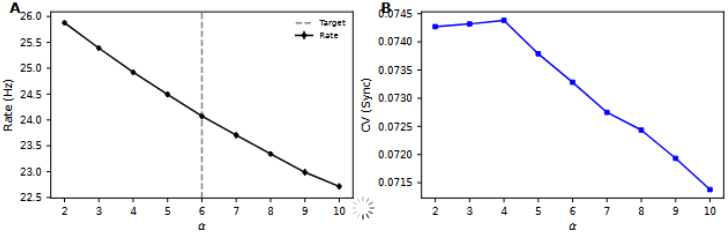
Calibration of global coupling scale *α*. (A) Mean population firing rate vs *α* (error bars: 95% CI from SEM across seeds). (B) Synchrony proxy (CV of smoothed population rate) vs *α*. Dashed line: chosen working point *α* = 6.

### 3.2 Strict attractor assay yields no evidence for bistability

Figure 4 illustrates the strict “kick-and-noise-off” assay. A strong transient pulse produces a temporary excursion, but the trajectory returns toward the control once the pulse ends. After noise is disabled, the trajectories do not remain separated; end-of-trial rates show no robust difference across seeds (paired Wilcoxon *p* = 9.91 *×* 10^−2^, *d*_*z*_ = − 0.25). Within this model class and operating regime, Sm dynamics appear monostable.

**Figure 4:**
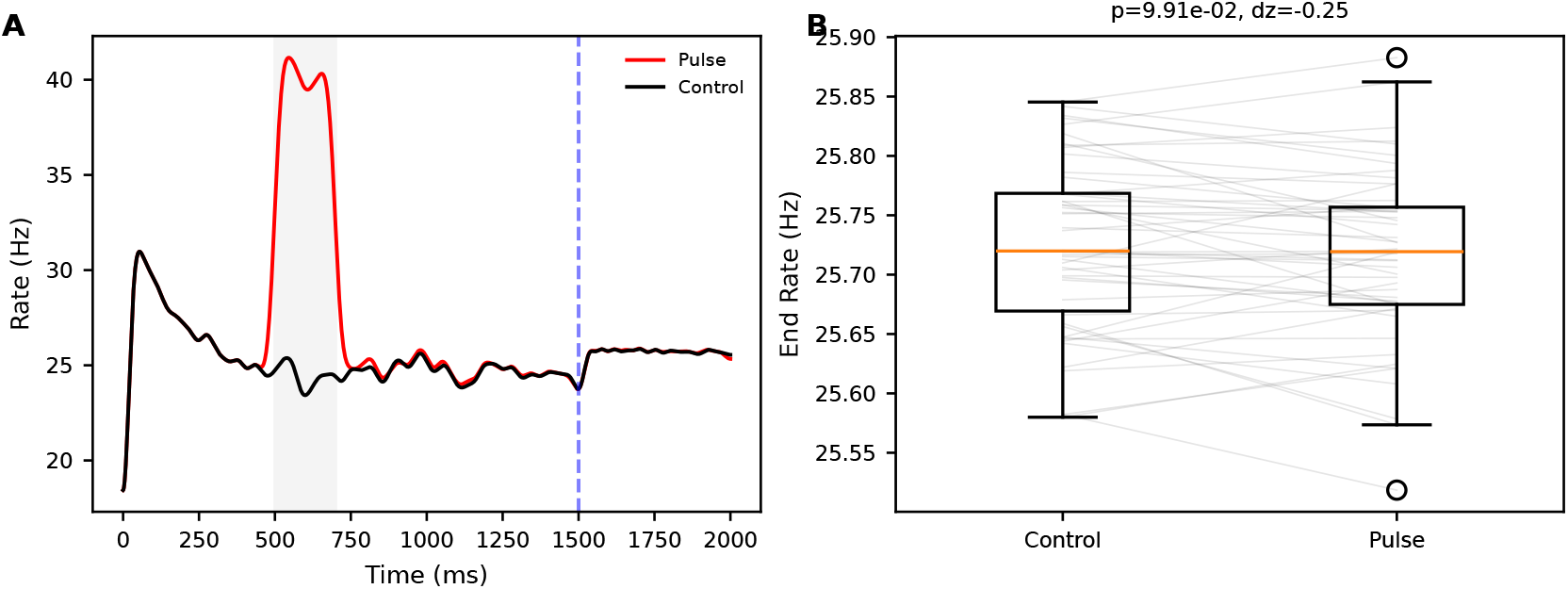
Strict test for bistable attractors. (A) Example population rate traces under Control (black) and Pulse (red). Dashed line: time when OU noise is turned off. (B) End-state firing rate (final window) across seeds shows no robust separation (paired Wilcoxon *p* = 9.91 *×* 10^−2^; within-subject *d*_*z*_ = −0.25).

A critical interpretation point: this negative result does *not* imply that Sm cannot support persistent internal states in vivo. It implies only that recurrence plus the present AdEx + current-based synapse instantiation is insufficient to generate a robust bistable attractor at the calibrated operating point.

### 3.3 Oscillations and feedback signatures

Population activity exhibits oscillatory structure in the population-rate signal. To assess whether 30–80 Hz band power depends on synaptically mediated interactions in this model, we compare: (i) a fully coupled simulation using the signed synapse-count adjacency *W* (Embedded; Full *W*), and (ii) an uncoupled control obtained by setting *W* = 0 (*W* = 0; labeled “Isolated” in Figure 5), which preserves single-neuron dynamics and injected OU noise but removes synaptic coupling.

**Figure 5:**
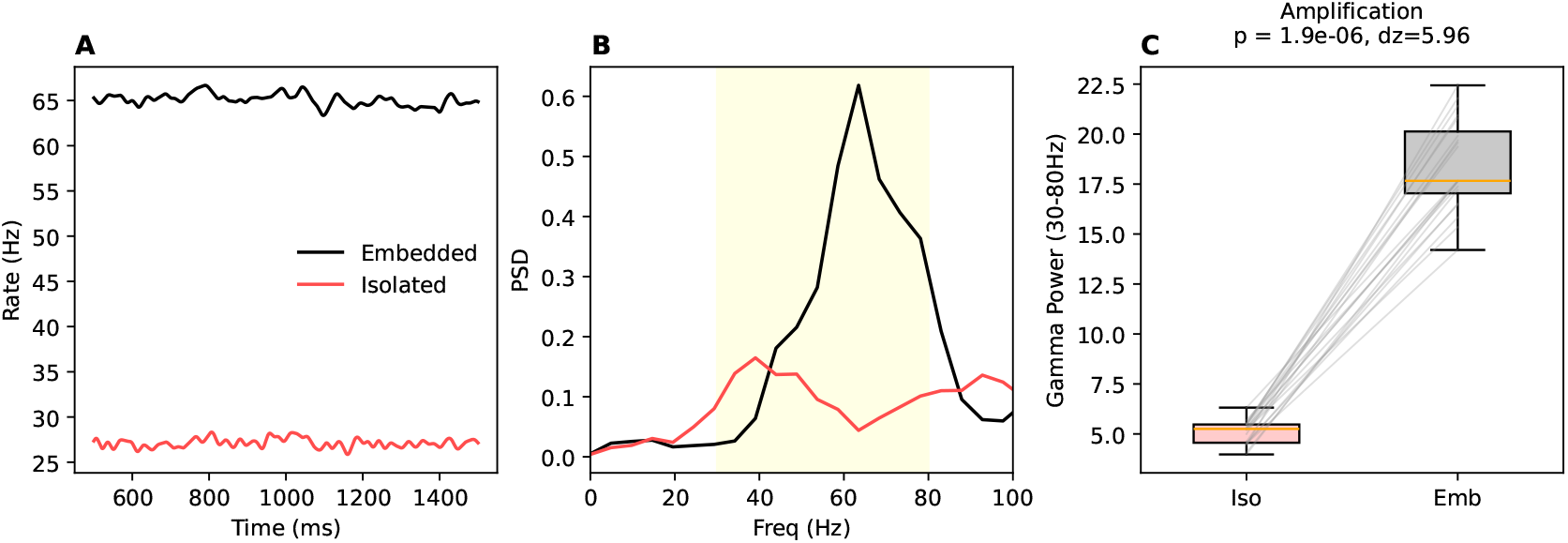
Synaptic coupling amplifies 30–80 Hz population-rate power. (A) Example population-rate traces for the fully coupled condition (Embedded; Full *W*) and the uncoupled control (*W* = 0, labeled “Isolated”). (B) PSD estimates (Welch) for the same traces; shaded band indicates 30–80 Hz. (C) Integrated 30–80 Hz power for the *W* = 0 control (Iso) and Full *W* (Emb) across *n* = 20 paired seeds (paired Wilcoxon signed-rank *p* = 1.9 × 10^−6^; *d*_*z*_ = 5.96).

**Figure 6:**
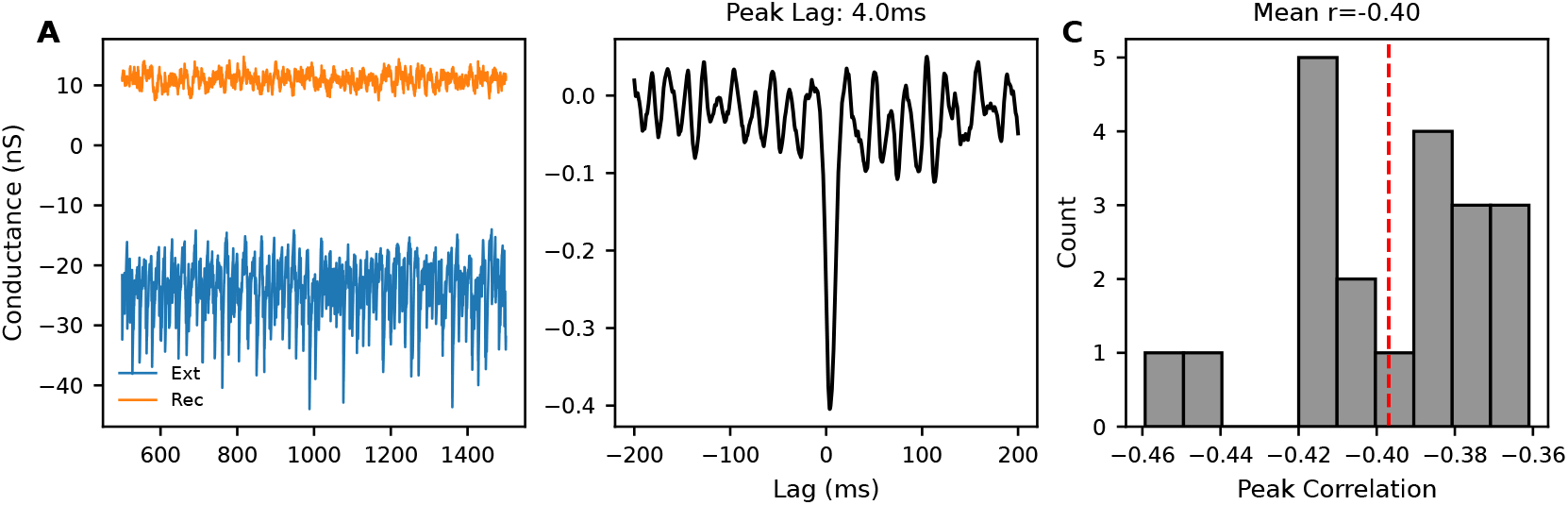
Input decomposition and fast negative coupling. (A) Decomposition of signed synaptic drive into external and recurrent components. (B) Normalized cross-correlation between population rate and external drive; strongest negative correlation occurs near zero lag (example peak lag ∼ 4 ms). (C) Distribution of peak negative correlation across seeds; mean peak correlation ≈ −0.40.

#### Synaptic coupling amplifies 30–80 Hz population-rate power

Across *n* = 20 paired random seeds, integrated 30–80 Hz power is substantially higher in the fully coupled condition:

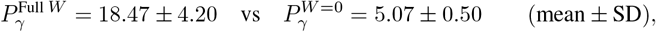

with a positive paired difference for all 20 seeds (Figure 5C). A paired Wilcoxon signed-rank test yields *p* = 1.9 *×* 10^−6^, with a large within-subject standardized paired effect size (*d*_*z*_ = 5.96).

#### Interpretation

This comparison establishes a strong amplification of population-rate spectral power in the 30–80 Hz band under synaptic coupling within the present AdEx + current-based synapse instantiation. Because coupling also alters mean population rate (Figure 5A), changes in 30–80 Hz power may reflect both rate-dependent spectral effects and coupling-dependent temporal structure. We therefore treat *P*_*γ*_ as a descriptive summary statistic rather than a mechanistic identifier of a specific oscillatory generator.

### 3.4 Negative controls and robustness probes

#### A stable linear dynamical baseline collapses to a fixed point

The linear baseline can track short-horizon fluctuations under teacher forcing, but fails in open-loop rollout, collapsing toward a trivial trajectory while the true rate remains oscillatory (Figure 7). This is the expected behavior of a stable linear system attempting to represent a limit cycle. The baseline is included as a cautionary control: local prediction accuracy does not guarantee correct long-horizon qualitative dynamics.

**Figure 7:**
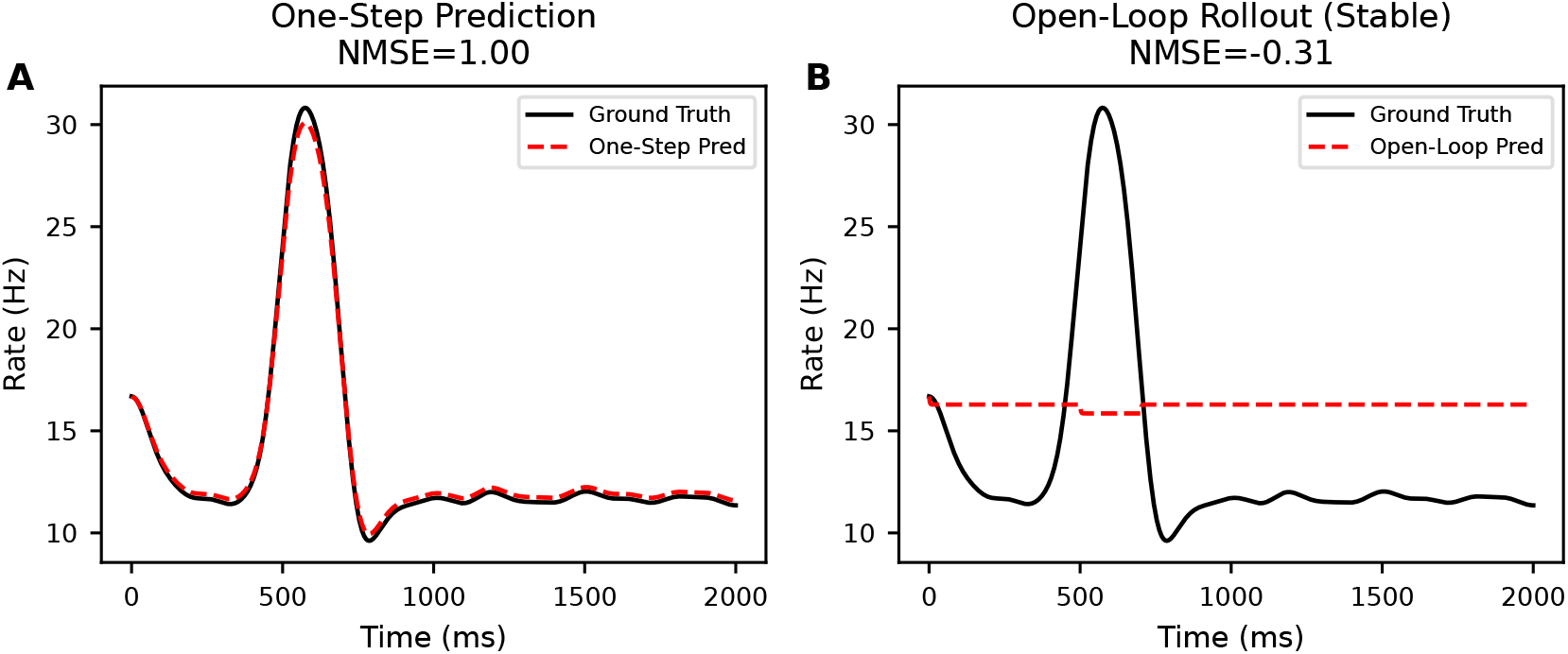
Linear baseline: one-step success, open-loop failure. (A) One-step prediction (teacher forced): the linear model tracks short-horizon fluctuations but exhibits bias. (B) Open-loop rollout: the stable linear model collapses toward a fixed point while the true trace remains oscillatory.

#### Robustness to neurotransmitter sign uncertainty (limited but nontrivial)

Finally, we perturb synaptic signs by randomly flipping a fraction of nonzero edges. Even with 30% edge sign flips, the mean absolute difference in population rate remains modest (mean 1.37 ± 0.01 Hz; Figure 8). This suggests that, for the present summary statistic (mean rate), the operating regime is not a knife-edge artifact of one exact sign assignment. We emphasize the qualifier: other observables (correlations, spectra, transient response) could be more sensitive.

**Figure 8:**
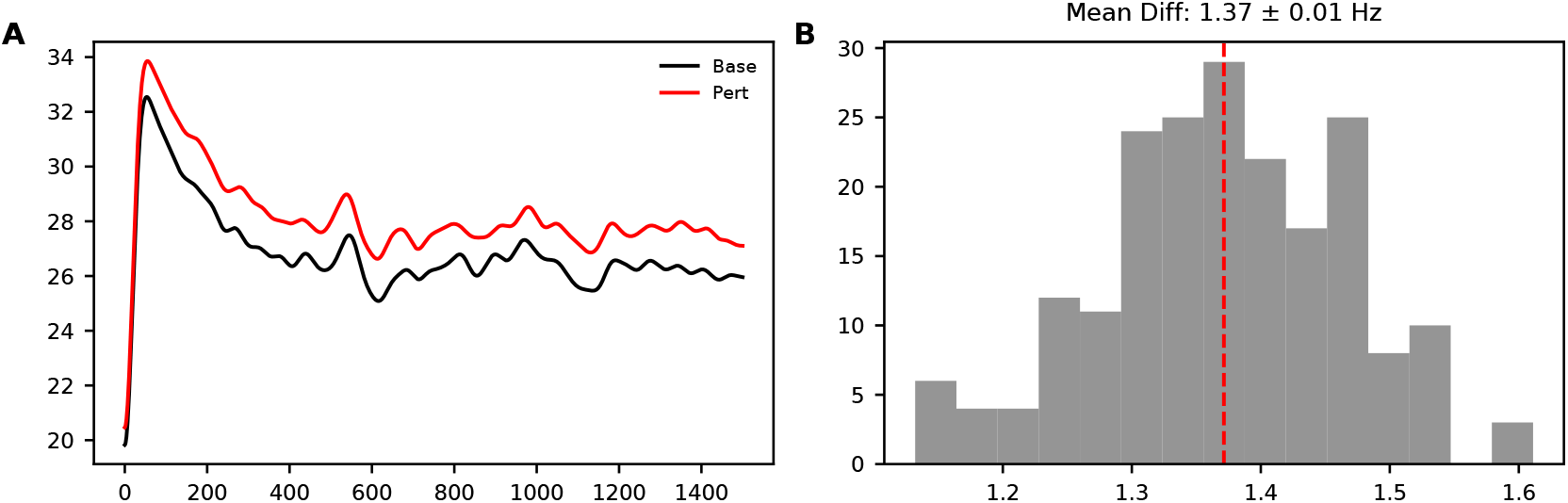
Sensitivity to synaptic sign perturbations. (A) Example population-rate traces under baseline signs and a 30% sign-flip perturbation. (B) Distribution of mean absolute rate differences across perturbations; mean difference 1.37 *±* 0.01 Hz (SEM).

## 4 Discussion

### 4.1 What we can (and cannot) conclude about Sm dynamics

#### No bistable “working memory” in this model class

The Sm family is strongly recurrent and anatomically positioned for temporal processing. Nevertheless, under a current-based AdEx instantiation calibrated to a stable non-saturating regime, we find no evidence for bistability under strict criteria. This is a meaningful negative result because it rules out a broad class of “recurrence implies memory” narratives *within* the tested assumptions. It does *not* rule out persistent states in vivo. Bistability can be introduced by mechanisms absent here: neuromodulatory state changes, short-term facilitation, NMDA-like slow synapses (not present in flies in the mammalian form), intrinsic bursting, structured sensory drive, or plasticity-induced changes.

#### Oscillations: synaptic coupling amplifies 30–80 Hz power, but interpretation should be modest

Gamma-band power is a widely used descriptive label, but “gamma power” can also reflect broadband changes in spiking and filtering. A useful warning from the broader oscillations literature is to distinguish true rhythmicity from generic increases in high-frequency power [11]. In our AdEx + current-based synapse instantiation, synaptic coupling produces a strong increase in integrated 30–80 Hz power relative to an uncoupled (*W* = 0) control, alongside an increase in mean population rate. We therefore treat *P*_*γ*_ as a descriptive summary statistic of high-frequency population-rate fluctuations rather than evidence for a specific oscillatory generator. A mechanistic interpretation would require controls that disentangle rate effects from rhythmic structure (e.g., rate-matched comparisons, peak-frequency stability, and coherence measures), as well as targeted in silico perturbations to identify which pathways or motifs drive the amplification.

#### Fast negative coupling suggests stabilizing feedback, not a full mechanism

Input decomposition reveals a short-lag negative correlation between external drive and the subcircuit rate. This is consistent with fast inhibitory feedback stabilizing the operating regime and shaping fluctuations. However, correlation is not causation: the effect could arise from shared drivers, delayed recurrence, or sign-policy artifacts. A mechanistic claim would require targeted in silico perturbations (e.g., removing specific external sources) and, ideally, experimental tests.

### 4.2 Why the “forensic” framing matters

Connectome-based modeling has a failure mode: seeing a pretty dynamical phenomenon once and calling it function. We attempted to make the opposite mistake: assume the phenomenon is an artifact until it survives multiple attempts to kill it. This stance leads naturally to: (1) explicit gates and seed replication, (2) negative controls that expose long-horizon errors, and (3) explicit stress tests for uncertain biological assumptions. The payoff is not a grand claim, but a reduction in self-deception.

### 4.3 Limitations and threats to validity

Several limitations are structural:

- **Model class:** current-based synapses and a homogeneous AdEx parameter set are a simplifying choice. Conductance-based effects and cell-type-specific excitability could change results.
- **Operating point:** *α* and *I*_bias_ are calibrated pragmatically; bistability might appear elsewhere in parameter space.
- **Sign mapping:** neurotransmitter predictions do not uniquely determine functional sign across all synapses; receptor expression and cotransmission matter.
- **Missing biology:** neuromodulation, gap junctions, and short-term plasticity are not modeled.
- **Observables:** we mainly analyze population rate. Subtype-resolved dynamics could show richer motifs.

We view these not as excuses, but as a map of what must be added before making stronger claims.

### 4.4 Practical next experiments

If one wanted to extend this work the most leverage-per-effort steps are: (i) a small sensitivity sweep over (*α, I*_bias_) for the attractor assay; (ii) adding modest heterogeneity (cell-type-specific excitability); and (iii) targeted in silico lesions of external sources to test the negative-coupling hypothesis.

## 5 Data and code availability

All code used for simulations, analyses, and figure generation is provided in the accompanying repository. Because FlyWire connectivity access may require credentials and may have redistribution restrictions, we do not assume raw adjacency matrices are universally shareable. We therefore provide scripts and provenance information (materialization identifiers and access dates) to re-fetch the Sm subgraph when permitted. A complete artifact bundle containing the manuscript sources and the exact code snapshot used to generate figures is included in the submission package.GitHub repository for the Sm-Forensic project

## A Forensic correctness gates

Our pipeline includes explicit checks to prevent silent failure modes. These gates are intentionally boring; they exist because “pretty figures” can be produced by broken pipelines.

1. **Nontriviality gate:** simulation must not be silent 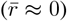 and must not saturate (implausibly high firing rates).
2. **Transfer sanity:** empirical *ϕ*(*I*) must be monotone and nonnegative; otherwise the neuron model or simulator is misconfigured.
3. **Seed replication:** each major claim is evaluated across multiple random seeds; single-seed “discoveries” are treated as anecdotes.
4. **Fixed analysis parameters:** PSD and filtering parameters are fixed across conditions to avoid p-hacking by changing preprocessing.
5. **Transient removal:** correlation analyses discard startup transients to reduce sensitivity to initial conditions.
6. ***p*-value range checks:** numerical outputs are validated to lie in legal ranges; a *p*-value of exactly 0 or 1 triggers a warning to inspect sensitivity and possible discretization/rounding artifacts.
7. **Negative controls:** linear dynamical baselines are run open-loop to demonstrate long-horizon qualitative mismatch.
8. **Assumption stress tests:** key uncertain assumptions (e.g., sign policy) are perturbed to assess regime robustness.

## B Model catalog and assumptions (clear-language summary)

This appendix enumerates the models used in the paper, what each is for, and what it is *not* for.

### B.1 Model 1: Connectome-derived weighted directed graph

**Object:** a directed weighted graph with synapse-count weights *W*_*ij*_. **Used for:** constraining which pairs of neurons can influence each other, and relative coupling strength. **Not claimed:** synapse counts are not absolute currents; they omit state dependence and receptor composition.

### B.2 Model 2: Spiking network instantiation (AdEx + exponential synapses)

**Object:** Eqs. (1)–(3) with OU noise Eq. (4). **Used for:** generating spike trains and population-rate dynamics under explicit assumptions. **Not claimed:** a biophysically fitted model of Sm cell types; we use a homogeneous parameterization as a transparent baseline.

### B.3 Model 3: Empirical transfer function *ϕ*(*I*)

**Object:** a measured input–output curve for the AdEx parameters. **Used for:** sanity checking monotonicity/saturation and providing a qualitative mean-field lens. **Not claimed:** a validated f-I curve for real Sm neurons.

### B.4 Model 4: Strict attractor assay

**Object:** a two-condition perturbation protocol with “noise off” after the kick. **Used for:** testing whether persistent separation exists in deterministic dynamics. **Not claimed:** a complete operationalization of memory; it is a minimal bistability test.

### B.5 Model 5: Linear dynamical baseline

**Object:** a 1D linear dynamical system Eq. (9) fit by ridge regression. **Used for:** showing that one-step prediction does not imply correct qualitative dynamics. **Not claimed:** a mechanistic model of Sm; it is a negative control.

### B.6 Model 6: Sign-perturbation stress test

**Object:** random perturbations of Σ by edge sign flips. **Used for:** evaluating robustness to neurotransmitter-to-sign uncertainty for coarse statistics. **Not claimed:** a faithful model of receptor expression; it is an uncertainty probe.

## C Mathematical and implementation details

### C.1 Exact discretization of OU noise

For Eq. (4), the exact update over a step Δ*t* is

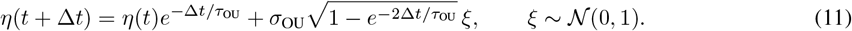

This avoids numerical drift associated with naive Euler–Maruyama discretization at small *τ*_OU_.

### C.2 Welch PSD and integrated band power

Welch’s method estimates *S*_*r*_(*f*) by averaging periodograms over overlapping windows. Integrated band power Eq. (7) is computed by numerical quadrature over the discrete frequency bins in the target band.

### C.3 Cross-correlation normalization

Given zero-mean signals *x*(*t*) and *y*(*t*), the normalized cross-correlation is

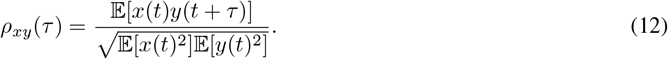

We report the most negative peak 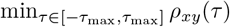 and its argmin.

### C.4 Ridge regression for the linear baseline

Let *x*_*t*_ be the population rate time series and define the design matrix *X* = [*x*_*t*_ 1] and target *y* = *x*_*t*+1_. Ridge regression solves

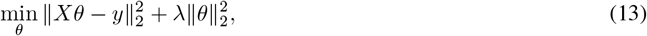

with closed-form solution 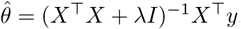. Open-loop rollout applies the learned mapping iteratively.

### C.5 Bootstrap confidence intervals

For descriptive statistics (e.g., mean peak correlation), we estimate confidence intervals by resampling seeds with replacement and recomputing the statistic.

## Notes

https://github.com/nalin-dhiman/sm-forensic

### Competing Interest Statement

The authors have declared no competing interest.

## References

[1] Sven Dorkenwald, Philipp Schlegel, et al. “A brain-wide map of neural circuitry in adult Drosophila”. In: Nature 634.8032 (2024), pp. 124–138.

[2] Arie Matsliah, Mehmet Fatih Keles, et al. “Neuronal parts list and wiring diagram for a visual system”. In: Nature 634.8032 (2024), pp. 166–180.

[3] Philipp Schlegel, Alexander S. Bates, et al. “Whole-brain annotation and multi-connectome cell typing quantifies circuit stereotypy in Drosophila”. In: Nature 634.8032 (2024), pp. 139–148.

[4] David A. Pospisil, Anindita Nandi, et al. “The fly connectome reveals a path to the effectome”. In: Nature 634.8032 (2024), pp. 201–209.

[5] Joshua R Sanes and S Lawrence Zipursky. “Design principles of insect and vertebrate visual systems”. In: Neuron 66.1 (2010), pp. 15–36. DOI: 10.1016/j.neuron.2010.01.018.

[6] Shin-ya Takemura et al. “Synaptic circuits and their variations within different columns in the visual system of Drosophila”. In: Proceedings of the National Academy of Sciences of the United States of America 112.44 (2015), pp. 13711–13716. DOI: 10.1073/pnas.1509820112.

[7] Romain Brette and Wulfram Gerstner. “Adaptive exponential integrate-and-fire model as an effective description of neuronal activity”. In: Journal of Neurophysiology 94.5 (2005), pp. 3637–3642.

[8] Wei Wei Liu and Rachel I. Wilson. “Glutamate is an inhibitory neurotransmitter in the Drosophila olfactory system”. In: Journal of Neuroscience 33.25 (2013), pp. 10245–10260.

[9] Frank G. Richter, Frank Mohrlen, et al. “Glutamate signaling in the fly visual system”. In: Frontiers in Cellular Neuroscience 12 (2018), p. 186.

[10] Susana Molina-Obando, Jairo F. Vargas-Fique, et al. “Visual system’s first synapse determines the sign of the response to light”. In: eLife 8 (2019), e43870.

[11] György Buzsáki and Xiao-Jing Wang. “Mechanisms of gamma oscillations”. In: Annual Review of Neuroscience 35 (2012), pp. 203–225. DOI: 10.1146/annurev-neuro-062111-150444.

